# Integrated Plasma and Tissue Proteomics Reveals Attractin Release by Intraluminal Thrombus of Abdominal Aortic Aneurysms and Improves Aneurysm Growth Prediction in Humans

**DOI:** 10.1101/2020.02.28.970483

**Authors:** Regent Lee, Ismail Cassimee, Honglei Huang, Pierfrancesco Lapolla, Anirudh Chandrashekar, Philip Charles, Benedikt Kessler, Roman Fischer, Ashok Handa, On behalf of the Oxford Abdominal Aortic Aneurysm Study

## Abstract

Background: Abdominal aortic aneurysms (AAA) are pathological dilatations of the aorta which can result in rupture and mortality. Novel methods of predicting AAA growth is a recognised priority in AAA research. Patient with AAAs have increased risk of cardiovascular morbidity. We have previously observed accelerated systemic endothelial dysfunction (measured by brachial artery FMD) in AAA patients and FMD correlates with future AAA growth. Further, systemic endothelial dysfunction is reversed by AAA repair. AAAs contain intra-luminal thrombus (ILT). Since ILT is either removed or excluded from circulation after successful repair of AAAs, we hypothesise that ILT to be the source of mediators that contribute to AAA growth. Methods: Patients were prospectively recruited to the Study (Ethics Ref SC/13/0250). Plasma samples were collected at baseline and at 1 year from each patient. Plasma samples were also collected before and at 10-12 weeks after surgery from each patient (n=29). Paired aneurysm wall, ILT, omental biopsies were collected intra-operatively during open surgical repair (n=3). In addition to analyses of the tissue, supernatant was obtained from ex vivo culture of these paired tissue samples. Samples were subjected to non-targeted LC-MSMS workflow after trypsin digest, using the Universal method to discover novel proteins. LC-MSMS data was analysed using the Progenesis QI pipeline. Results: The median AAA size at baseline was 48 mm. 59 patients were prospectively followed for 12 months. The median growth rate of AAA was 3.8%/year (IQR 1.9% to 6.8%). Comparison between patients with the fastest vs the slowest (n=10 each) showed 116 proteins to be differentially expressed in their plasma. Among these proteins, 35 also changed significantly before and after AAA repair, suggesting their origin to from the AAA complex. Comparison of the proteomics profile of aneurysm tissue, ILT, and omental artery show 128 proteins to be uniquely present in ILT. Analyses of the tissue culture supernatant further revealed 3 proteins that are: (i) uniquely present in ILT; (ii) released by ILT; (iii) systemic levels reduced after AAA surgery; (iv) differs between fast and slow growth AAAs. One of these proteins is attractin. To validate the LC-MSMS data, attractin level in individual patient was measured by ELISA. Consistent with the LC-MSMS data, plasma attractin level is higher in patients with fast AAA growth. Plasma attractin level correlates significantly with future AAA growth rate (Spearman r=0.35, P<;0.005). Using attractin and AAA diameter as input variables, the AUROC for predicting no growth of AAA at 12 months is 85% (P<0.001). Conclusion: We show that ILT of AAAs releases mediators (such as attractin) during the natural history of AAA growth. These are novel biomarkers for AAA growth prediction in humans.

---

Abdominal aortic aneurysms (AAAs) are pathological dilatation of the abdominal aorta to larger than 30mm in diameter. Left untreated, it eventually results in AAA rupture and high mortality. Methods for the prediction of AAA growth is considered as a priority for research in the opinions of our peers^1^. It can guide different aspects of clinical management in terms of the frequency of monitoring of AAAs and the optimal timing for surgery.

The Oxford Abdominal Aortic Aneurysm (OxAAA) Study has previously reported a method of AAA growth prediction by incorporating 9 circulating proteins (derived using a commercially available protein array), flow mediated dilatation of brachial artery (FMD, a physiological marker of systemic endothelial function), and AAA diameter^2^. We had also observed that systemic endothelial function (measured by brachial artery FMD) deteriorates during the natural history of AAA growth and is reversed by AAA repair^3^.

Most AAAs contain intra-luminal thrombus (ILT)^4^. Since ILT is either removed or excluded from circulation after successful repair of AAAs, we hypothesise that ILT is the source of mediators that contribute to AAA growth. In this report, we utilised mass spectrometry analyses on blood, thrombus tissue, and tissue supernatant collected from patients during the natural history of AAA progression to discover novel predictors of AAA growth in humans.

Details regarding the OxAAA study cohort and recruitment process have been published^3^. In brief, this prospective study (Ethics Ref: 13/SC/0250) recruited patients in the National Health Service setting. Baseline assessments were performed. In addition to the measurement of AAA antero-posterior diameter by ultrasound imaging, fasting blood sample was collected and Platelet-poor plasma was prepared using two-staged centrifugation as previously described^5^ and stored at −80□C for subsequent analysis. Prospective AAA annual growth rates were calculated based on the diameter measurements in the subsequent AAA monitoring ultrasound scans.

Plasma samples were collected at baseline and at 1 year from each patient (n=59). Based on the prospectively recorded aneurysm growth rates, we selected a subset of patients [fastest (n=10) vs slowest (n=10)]. These were pooled for the initial discovery analysis. Plasma samples were also collected before and at 10-12 weeks after surgery from each patient (n=29). *Paired* aneurysm wall, ILT, omental biopsies were collected intra-operatively during open surgical repair (n=3). In addition to analyses of the tissue, supernatant was obtained from *ex vivo* culture of these paired tissue samples. We utilised a similar approach for plasma biomarker discovery as recently described^6^. Samples were subjected to Liquid Chromatography Tandem Mass Spectrometry (LC-MS/MS) proteomic analysis to discover protein level differences^6^. LC-MS/MS data were analysed using the Progenesis QI software (NonLinear Dynamics), and included only proteins with at least two matched peptide sequences.

The median AAA size at baseline was 48 mm. The median growth rate of AAA was 3.8%/year (IQR 1.9% to 6.8%). Comparison between patients with the fastest vs the slowest AAA growth showed 116 proteins (listed by the UniProt Protein IDs in Figure Panel 1) to be differentially expressed in their plasma (Figure panel 2-A). Among these proteins, 35 also changed significantly before and after AAA repair (Figure panel 2-B), suggesting their origin from the AAA complex. Comparison of the proteomics profile of aneurysm tissue, ILT, and omental artery show 128 proteins to be uniquely present in ILT (Figure panel 2-C).

**Figure:**
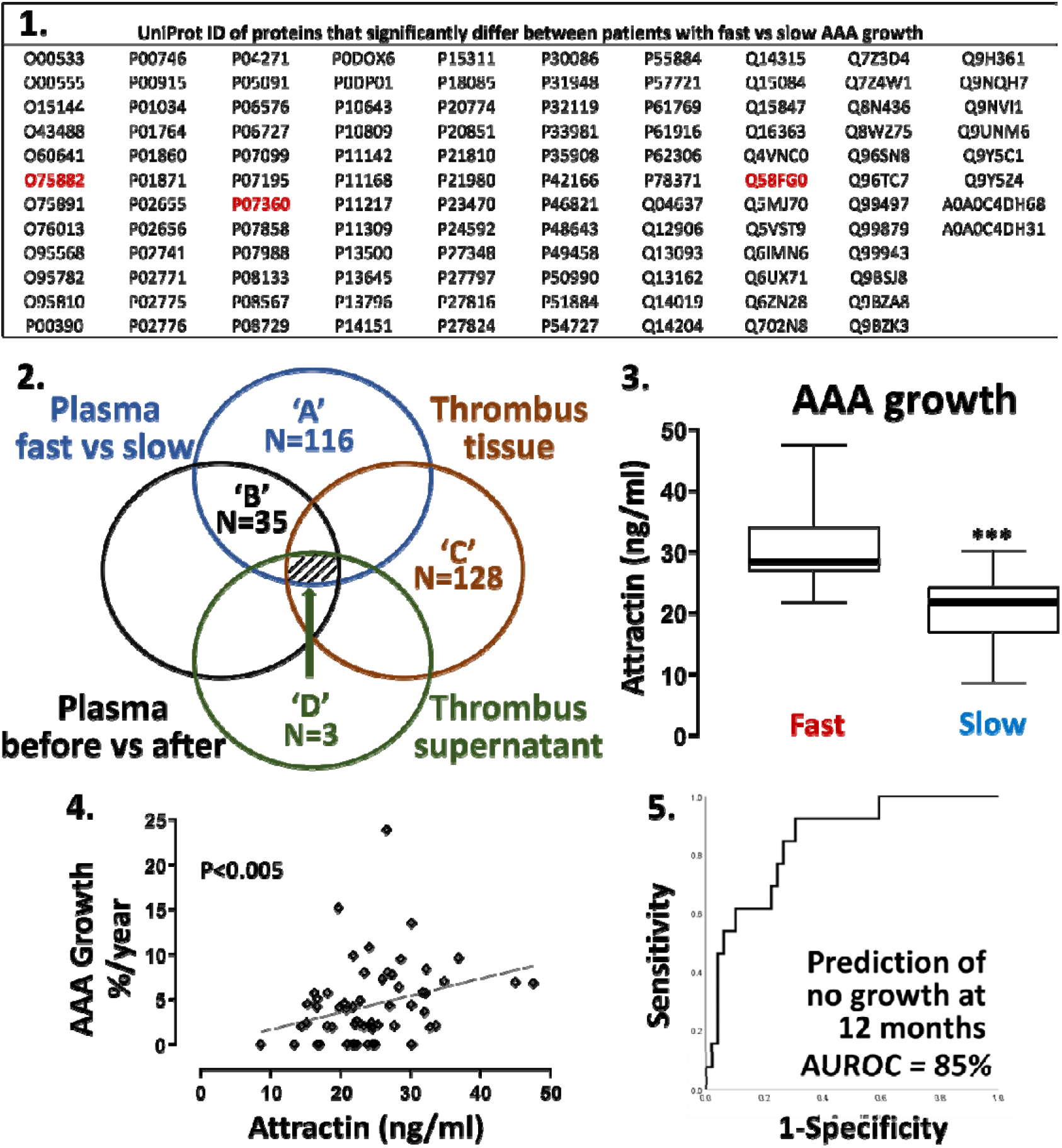
Integrated Plasma and Tissue Proteomics Reveals Attractin Release by Intraluminal Thrombus of Abdominal Aortic Aneurysms and Improves Aneurysm Growth Prediction in Humans. AAA growth rates were prospectively recorded in 59 patients. Based on the growth rate in the subsequent 12 months, we selected a subset of patients (fastest vs slowest, n=10 each) for the initial discovery analysis. Plasma samples were also collected from patients before and after AAA repair (n=29). Paired aneurysm wall, ILT, omental biopsies were collected intra-operatively during open surgical repair (n=3). In addition to analyses of the tissue, supernatant was obtained from ex *vivo* culture of these paired tissue samples. Samples were subjected to Liquid Chromatography Tandem Mass Spectrometry (LC-MS/MS) proteomic analysis to discover protein level differences. Plasma samples were pooled in their respective groups for analysis. Other samples were analysed individually. LC-MS/MS data were analysed using the Progenesis QI software (NonLinear Dynamics) and included only proteins with at least two matched peptide sequences. Comparison between patients with the fastest vs the slowest aneurysm growth showed 116 proteins (listed by the UniProt Protein IDs in Panel 1) to be differentially expressed in their plasma (2-A). Among these proteins, 35 also changed significantly before and after AAA repair (2-B), suggesting their origin from the AAA complex. Comparison of the proteomics profile of aneurysm tissue, ILT, and omental artery showed 128 proteins to be uniquely present in ILT (2-C). Analyses of the tissue culture supernatant further revealed 3 proteins that were: (i) uniquely present in ILT; (ii) released by ILT; (iii) systemic levels reduced after AAA surgery; (iv) different between fast and slow growth AAAs (2-D). These are: attractin (UniProt ID O75882), complement C8 (UniProt ID P07360), heat shock protein AA5P (HSPAA5P, UniProt ID Q58FG0). Attractin level in individual patient was further measured by ELISA (R&D Quantikine DATRN0). Plasma attractin level is significantly higher in patients with fast AAA growth (panel 3, median 28.5 vs 21.9 ng/ml, P<0.001). Plasma attractin level correlates significantly with future AAA growth rate (Figure panel 4, Spearman r=0.35, P<0.005). We regressed the measured values of attractin in combination with AAA diameter against a categorical response with levels of ‘Slow/no’ growth (0%) or growth (>0% growth) for outcomes at 12 months. Using attractin and AAA diameter as input variables, the AUROC for predicting no growth of AAA at 12 months is 85% (panel 5, asymptotic P<0.001).

Analyses of the tissue culture supernatant further revealed 3 proteins that were: (i) uniquely present in ILT; (ii) released by ILT; (iii) reduced in systemic circulation after AAA surgery; (iv)different between fast and slow growth AAAs (Figure panel 2-D). These are: attractin (UniProt ID O75882), complement C8 (UniProt ID P07360), heat shock protein AA5P (UniProt ID Q58FG0). To technically validate the LC-MS/MS data, attractin level in individual patient was measured by ELISA (R&D Quantikine DATRN0). Consistent with the LC-MS/MS data, plasma attractin level is significantly higher in patients with fast AAA growth (Figure panel 3, median 28.5 vs 21.9 ng/ml, P<0.001). Plasma attractin level correlates significantly with future AAA growth rate (Figure panel 4, Spearman r=0.35, P<0.005).

We tested the utility of a generalised linear model to predict aneurysm growth in these patients. We regressed the measured values of attractin in combination with the measurements of AAA diameter against a categorical response with levels of ‘Slow/no’ growth (0%) or growth (>0% growth) for outcomes at 12 months. Using attractin and AAA diameter as input variables, the area under receiver operating characteristics (AUROC) for predicting no growth of AAA at 12 months is 85% (Figure panel 5, asymptotic P<0.001) as compared to 76% with AAA diameter alone.

This report is a significant breakthrough from our previous work^2^. By focusing on the role of thrombus as a source of systemic mediator release, we discover novel proteins that are released from thrombus and drive AAA growth in humans, tested in a prospectively recruited cohort. This is the first report in which novel proteins that correlate to future AAA growth have been discovered through a mass spectrometry workflow. The validity of the mass spectrometry discovery workflow is demonstrated by the precise replication of the LC-MS/MS data by ELISA measurements on individual patient samples.

Since the first description of attractin in 1998^7^, there has been little in the reported literature regarding its biological role in disease. Evidence points toward its release by activated T-cell. It is involved in the initial immune cell clustering during inflammatory response and it regulates chemotactic activity of chemokines^7^. There is mounting evidence of T-cells being active in AAA ILT and that it plays a role in AAA pathophysiology^8^. This report is the first to implicate attractin in human AAA progression and warrants further mechanistic investigations.

It is important for external cohorts to replicate the efficacy of our biomarker panel. We hope this work serves as a primer to generate interest in the vascular surgical community and stimulates future efforts to validate the prediction algorithm.

## Acknowledgement

The Oxford Abdominal Aortic Aneurysm Study was supported by the following: University of Oxford, Medical Sciences Division Medical Research Fund (MRF/HT2016/2191); University of Oxford, Nuffield Department of Surgical Sciences; John Fell Oxford University Press Research Fund (142/075); National Institute of Health Research (NIHR) Oxford Biomedical Research Centre; British Heart Foundation Centre of Research Excellence, Oxford (RE/13/1/30181); RL was supported by a Academy of Medical Science Starter Grant, UK (SGL013/1015). PL was supported by an EU Erasmus÷ traineeship studentship. BMK and PDC were supported by a John Fell Oxford University Press Research Fund (133/075) and a Welcome Trust Grant (097813/11/Z).

## References

1. Lee R, Jones A, Cassimjee I and Handa A. International opinion on priorities in research for small abdominal aortic aneurysms and the potential path for research to impact clinical management. Int J Cardiol. 2017;245:253–255.

2. Lee R, Charles PD, Lapolla P, Cassimjee I, Huang H, Kessler B, Fischer R and Handa A. Integrated Physiological and Biochemical Assessments for the Prediction of Growth of Abdominal Aortic Aneurysms in Humans. Ann Surg. 2018.

3. Lee R, Bellamkonda K, Jones A, Killough N, Woodgate F, Williams M, Cassimjee I and Handa A. Flow Mediated Dilatation and Progression of Abdominal Aortic Aneurysms. Eur J Vasc Endovasc Surg. 2017;53:820–829.

4. Whaley ZL, Cassimjee I, Novak Z, Rowland D, Lapolla P, Chandrashekar A, Pearce BJ, Beck AW, Handa A and Lee R. The Spatial Morphology of Intraluminal Thrombus Influences Type II Endoleak After Endovascular Repair of Abdominal Aortic Aneurysms. Ann Vasc Surg. 2019.

5. Lee R, Antonopoulos AS, Alexopoulou Z, Margaritis M, Kharbanda RK, Choudhury RP, Antoniades C and Channon KM. Artifactual elevation of plasma sCD40L by residual platelets in patients with coronary artery disease. Int J Cardiol. 2013;168:1648–50.

6. Lee R, Fischer R, Charles PD, Adlam D, Valli A, Di Gleria K, Kharbanda RK, Choudhury RP, Antoniades C, Kessler BM and Channon KM. A novel workflow combining plaque imaging, plaque and plasma proteomics identifies biomarkers of human coronary atherosclerotic plaque disruption. Clin Proteomics. 2017; 14:22.

7. Duke-Cohan JS, Tang W and Schlossman SF. Attractin: a cub-family protease involved in T cell-monocyte/macrophage interactions. Adv Exp Med Biol. 2000;477:173–85.

8. Ismail Cassimjee RLaJP. Inflammatory Mediators in Abdominal Aortic Aneurysms. 2017.

